# Prevalence and Associated Risk Factors of Bovine Fasciolosis in Bahir Dar, Ethiopia: Cross-Sectional Study

**DOI:** 10.1101/2024.08.07.606800

**Authors:** Tesfaye Mesfin, Theobesta Solomon, Abraham Belete Temesgen

## Abstract

Cattle are among the most important livestock resources in Ethiopia, contributing significantly to the agricultural economy and rural livelihoods. They provide meat, milk, hides, draft power for crop production, and serve as a major source of income for farmers. Despite their vital role, cattle productivity is often constrained by various diseases, particularly parasitic diseases. One of the most significant of these is bovine fasciolosis, a condition caused by ingestion of metacercariae of liver flukes belonging to the genus *Fasciola*. This study aimed to assess the prevalence and associated risk factors of bovine fasciolosis in bahir dar, Ethiopia. A cross-sectional study was conducted from November 2021 to April 2022. A total of 384 cattle were randomly selected from different locations within the study area. Animals of all age groups and both sexes were included. Fecal samples were collected directly from the rectum of each animal using clean, labeled containers. The samples were examined using standard coprological techniques, specifically the sedimentation method, to detect liver fluke eggs. All findings were recorded, and the data were analyzed using descriptive statistical methods. The overall prevalence of fasciolosis was 49.21% (n=189). Based on origin, Sebatamit had the most incidence at 61.84% (n=47), followed by Kebele 11 at 59.37% (n=57), Tikurit at 50% (n=59), and Latammba at 27.65% (n=26). Statistical analysis revealed significant disparities in occurrence among areas. Cattle in poor condition had the largest prevalence (n=80, 64%), followed by medium condition (n=85, 50%) and fat cattle (n=24, 26.96%). This variation was statistically significant. Age-group analysis revealed comparable prevalence rates, with young cattle at 50.38% (n=65), adults at 47.33% (n=71), and elderly cattle at 50.47% (n=53), with no significant differences found. There were no significant sex-related variations in prevalence, with males exhibiting a prevalence of 49.73% (n=93) and females 48.73% (n=96). Local cattle had a slightly higher prevalence (n=111, 51.62%) than crossbreds (n=78, 46.15%), although the difference was not statistically significant (P=.29).These findings underscore the need for targeted, location-specific control strategies and highlight the importance of improved nutritional and health management practices to reduce the burden of fasciolosis in cattle populations.

## 1. Introduction

Ethiopia hosts one of the largest livestock populations in Africa, with an estimated 55.03 million cattle, 27.32 million sheep, and 28.16 million goats as of 2019. Cattle, in particular, play a central role in the country’s agricultural economy, with the dairy sector contributing over 81% of total milk production. Despite this abundance, livestock productivity remains low due to constraints such as poor nutrition, inadequate management, and widespread infectious diseases. Among these, fasciolosis is one of the most impactful parasitic diseases, causing substantial economic losses through reduced growth, impaired fertility, decreased milk yield, and increased mortality [1].

Fasciolosis is caused by trematode parasites of the genus *Fasciola*, commonly known as liver flukes. Infection occurs when animals ingest metacercariae, the infective stage of the parasite, from contaminated pasture, water, or feed. Two species are primarily responsible: *Fasciola hepatica*, which predominates in temperate regions, and *Fasciola gigantica*, more common in tropical climates, including much of Africa and Ethiopia [2–4]. Transmission relies on the presence of aquatic snails, such as *Lymnaea truncatula* and *Lymnaea natalensis*, which serve as intermediate hosts under suitable conditions like stagnant water and moist environments [5].

Infected cattle experience liver tissue damage due to migrating immature flukes, resulting in inflammation, bile duct obstruction, hepatocellular necrosis, and fibrosis. Clinical signs often include weight loss, jaundice, poor body condition, and reduced productivity. Severe infections compromise liver function, predispose animals to secondary infections, and may lead to significant morbidity and mortality [6]. Economically, fasciolosis causes extensive liver condemnation at slaughterhouses, higher veterinary costs, and financial losses for farmers and the meat industry [7].

Epidemiology of fasciolosis is influenced by host and environmental factors, including age, sex, breed, management practices, and ecological conditions. Older animals are often more affected due to cumulative exposure, while some breeds show variable susceptibility. Differences in grazing behavior and reproductive cycles may contribute to higher prevalence in females in some cases [8–10].

Additionally, pasture type, access to contaminated water, and use of anthelmintics play critical roles in transmission. Although previous studies have provided insights into fasciolosis in Ethiopia, many were localized, leaving gaps in regional prevalence and risk factors. Given the ecological and management diversity across the country, region-specific studies are essential to inform targeted control strategies. In particular, the central and northern highlands, where cattle production is economically significant, may present unique environmental conditions that affect parasite dynamics [11–14]. Therefore, this study aimed to determine the prevalence and associated risk factors of bovine fasciolosis in bahir dar, Ethiopia.

## 2. Materials and Methods

### 2.1. Study Area

The study was conducted in Bahir Dar, Ethiopia, from November 2021 to April 2022 (Figure 1). Bahir Dar is located approximately 575 km northwest of Addis Ababa, at an elevation of 1500–2600 meters above sea level, with geographic coordinates of 12°29′ N latitude and 37°29′ E longitude. The area receives an average annual rainfall of 1200–1600 mm, and temperatures range from 8 to 31 °C. The landscape is predominantly plain plateaus, covering roughly 70% of the region, and the vegetation includes shrub formations, low woodlands, evergreen areas, and semi-humid highland vegetation. Agriculture is a key livelihood, with major crops including teff (*Eragrostis tef*), wheat (*Triticum aestivum*), maize (*Zea mays*), and various pulses [1].

**Figure 1.**
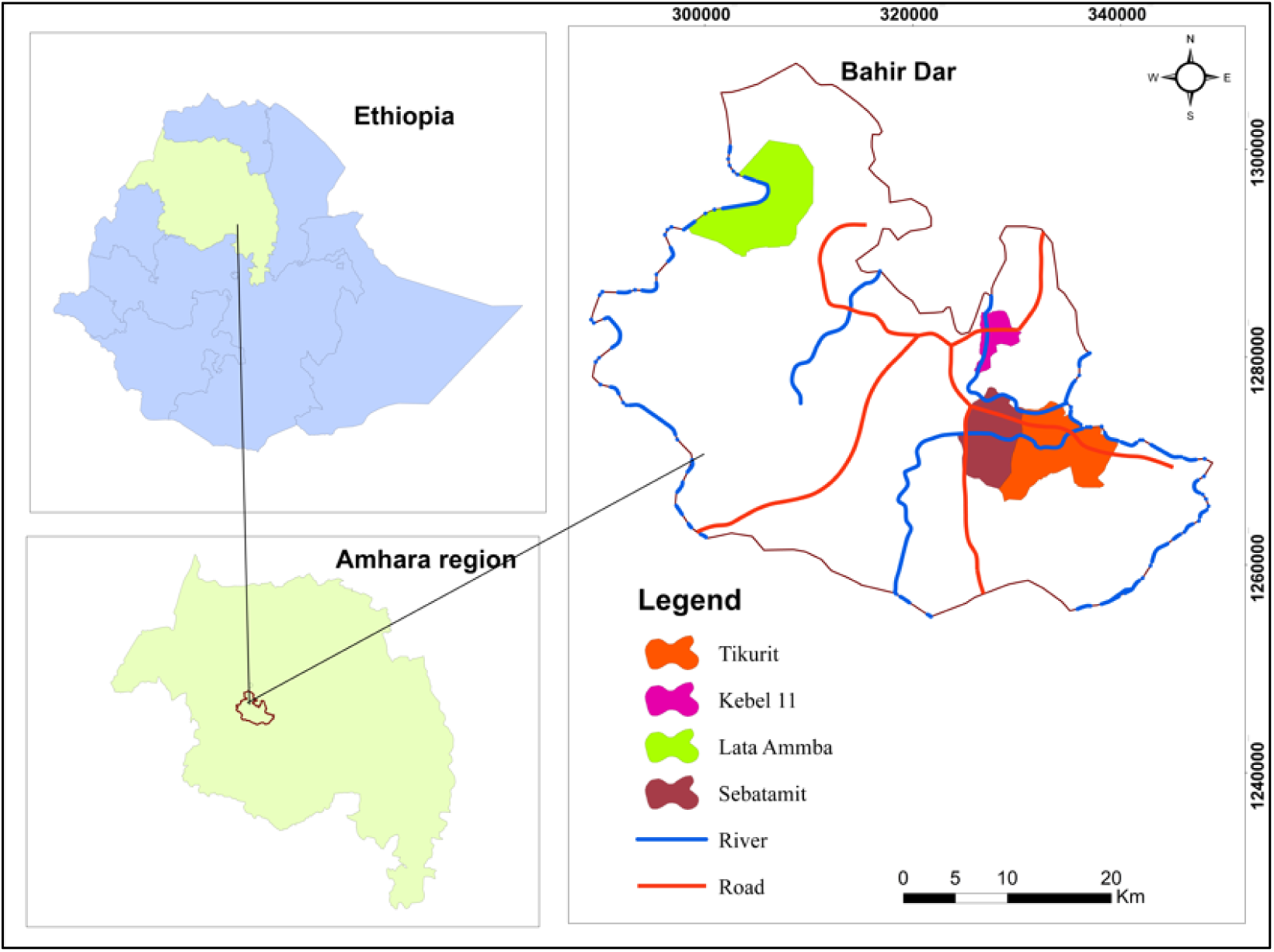
Map of the Study Area (Arc GIS 10.8)

### 2.2. Study Animal and Sampling Method

The study animals consisted of cattle from four selected sites in Bahir Dar: Kebele 11, Sebatamit, Tikurit, and Latammba. Both indigenous and crossbred Holstein Friesian cattle reared under local management conditions were included. A total of 384 cattle were randomly selected, representing both sexes and multiple age groups. Animal age was assessed based on dentition and categorized as young or adult [15]. Body condition score (BCS) was also evaluated for each animal following established guidelines to estimate nutritional and health status [16].

### 2.3. Study Design and Sample Size

A cross-sectional study was carried out in Bahir Dar, Ethiopia, from November 2021 to April 2022 to determine the prevalence and associated risk factors of bovine fasciolosis in bahir dar, Ethiopia. The risk factors considered included origin (location), breed, age, sex, and body condition of the cattle. The required sample size for fecal sample collection was calculated using a 95% confidence level, 5% absolute precision, and an expected prevalence of 50% in the absence of prior data for the study area [17].

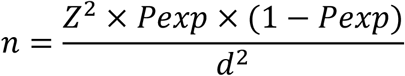

Where: *n* = required sample size; *P*exp = expected prevalence (0.5); *d* = desired absolute precision (0.05); *Z* = Z-value for confidence level (1.96). Based on this formula, a total of 384 cattle were included in the study.

### 2.4. Fecal Examination and Identification of Fasciola Eggs

Fresh fecal samples were collected directly from the rectum of each animal using disposable gloves and placed into universal bottles containing 10% formalin as a preservative. Samples were transported under controlled conditions to the Bahir Dar Regional Veterinary Laboratory for parasitological examination. The sedimentation technique was used to detect Fasciola eggs, following standard procedures [18,19]. To differentiate Fasciola eggs from those of other trematodes, such as Paramphistomum species, the sediment was stained with methylene blue. Fasciola eggs are typically yellowish, large, and operculated, whereas Paramphistomum eggs stain blue [20,21].

### 2.5. Data Management and Analysis

The raw data collected from the study were organized and entered into a Microsoft Excel spreadsheet for initial management. Subsequently, the data were exported to STATA version 16.0 (StataCorp, College Station, TX, USA) for statistical analysis. A chi-square (χ^2^) test was used to evaluate the correlation between infection rates and risk factors such as age, sex, breed, and location. The test evaluated infection rates based on these parameters, with a significance level of P < 0.05.

## 3. Results

### 3.1. Prevalence of Bovine Fasciolosis

The overall prevalence of bovine fasciolosis was 49.21% (n = 189). Based on origin, Sebatamit showed the highest prevalence at 61.84% (n = 47), followed by Kebele 11 at 59.37% (n = 57), Tikurit at 50% (n = 59), and Latammba at 27.65% (n = 26). Statistical analysis indicated a significant variation in prevalence among the study sites. Body condition also showed marked differences, with poor-condition cattle exhibiting the highest prevalence (n = 80, 64%), followed by medium-condition animals (n = 85, 50%), and fat cattle (n = 24, 26.96%); this variation was statistically significant. Age-related prevalence was comparable, with young cattle at 50.38% (n = 65), adults at 47.33% (n = 71), and older cattle at 50.47% (n = 53), showing no significant difference. Sex did not influence infection rates, with prevalences of 49.73% (n = 93) in males and 48.73% (n = 96) in females. Similarly, breed showed no significant effect, although local cattle had a slightly higher prevalence (n = 111, 51.62%) compared to crossbreds (n = 78, 46.15%), with no significant difference (*P* = .29) (Table 1).

**Table 1.** Prevalence of bovine fasciolosis based on risk factors.

| Risk factor and category | Examined (n=384) | Positive (n=189), % | Chi-square (*df*) | *P* value |
| --- | --- | --- | --- | --- |
| Origin |  |  |  |  |
| Tikurit | 118 | 59 (50) | 26.31 (3) | <.001 |
| Sebatamit | 76 | 47 (61.84) |  |  |
| Latammba | 94 | 26 (27.65) |  |  |
| Kebele 11 | 96 | 57 (59.37) |  |  |
| Body condition |  |  |  |  |
| Poor | 125 | 80 (64) | 28.60 (2) | <.001 |
| Medium | 170 | 85 (50) |  |  |
| Fat | 89 | 24 (26.96) |  |  |
| Age |  |  |  |  |
| Young | 129 | 65 (50.38) | 0.839 (2) | .35 |
| Adult | 150 | 71 (47.33) |  |  |
| Old | 105 | 53 (50.47) |  |  |
| Sex |  |  |  |  |
| Male | 187 | 93 (49.73) | 0.844 (1) | .35 |
| Female | 197 | 96 (48.73) |  |  |
| Breed |  |  |  |  |
| Local | 215 | 111 (51.62) | 1.134 (1) | .29 |
| Crossbred | 169 | 78 (46.15) |  |  |

### 3.2. Identification of Fasciola Eggs

The Fasciola eggs detected in cattle fecal samples are shown in Figure 2. Identification was based on morphological characteristics [20,21]. The eggs are ovoid, possess a thick yellowish-brown shell, and typically have an operculum at one end.

**Figure 2.**
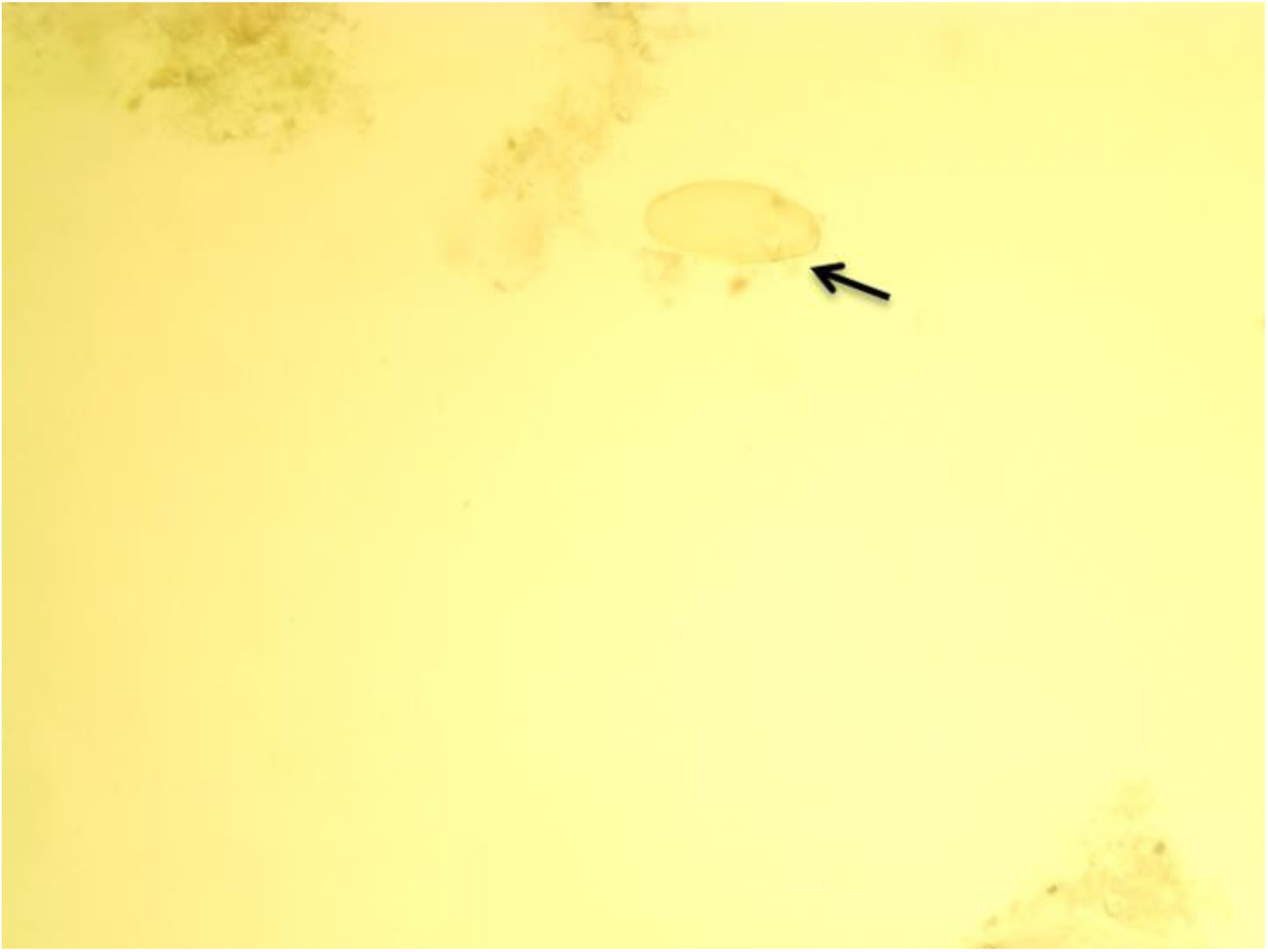
Egg of Fasciola species: yellowish egg (arrow).

## 4. Discussion

Bovine fasciolosis remains one of the most important parasitic diseases affecting cattle in Ethiopia, causing substantial economic losses through reduced growth, poor milk production, liver condemnation, and increased susceptibility to secondary infections. The present study recorded an overall prevalence of bovine fasciolosis of 49.21% in the study area, which aligns with previous reports from different regions of Ethiopia. For instance, prevalence rates of 41.41% and 54.5% have been reported in Woreta [22] and Jimma [23], respectively. Similar rates were observed in North-East Amhara (47.10%) [13], and Eastern Shoa, Kuyu District (54.2%) [19], likely associated with common agroecological factors such as the presence of *Lymnaea* snails and husbandry practices like communal water use and irrigation that facilitate pasture contamination [24].

Conversely, lower prevalence rates have been reported in certain areas. Studies in Soddo (4.9%) [25], Nekemte (15.9%) [7], and Southern Ethiopia (15.9%) [8] demonstrated lower frequencies, while Zenzelma, Bahir Dar (26%) [1], and Bahir Dar (32.3%) [26] reported slightly lower prevalence. These variations are likely influenced by differences in altitude, geography, climate, snail host abundance, management practices, and anthelmintic use [27].

Prevalence also varied significantly among study sites in the current work, with Sebatamit recording the highest prevalence (61.84%), followed by Kebele 11 (59.37%), Tikurit (50%), and Latammba (27.65%). This observation is consistent with prior studies indicating substantial geographical variation in fasciolosis prevalence [28–31], which may be influenced by climatic conditions, animal management practices, and access to veterinary services [32].

Cattle in poor body condition exhibited the highest fasciolosis prevalence (64%), compared to medium (50%) and fat cattle (26.96%). This pattern corresponds with several Ethiopian reports that found significantly higher infection rates in animals with poor body condition, likely reflecting increased susceptibility due to malnutrition, compromised immunity, and concurrent infections that weaken host defenses [33–36]. Poor body condition may also result from chronic disease or environmental stressors that predispose cattle to parasitic infection. However, some studies reported no significant differences in prevalence among body condition categories, indicating that body condition may not always predict *Fasciola* infection [23,37].

In this study, no significant differences were observed across age groups, with young cattle (50.38%), adults (47.33%), and older cattle (50.47%) showing similar prevalence. Several studies in Ethiopia have likewise reported no significant association between age and *Fasciola* infection [38,39], although other reports documented age-related variation, likely reflecting differences in exposure or management practices [40,41].

Similarly, sex did not significantly influence prevalence, with males at 49.73% and females at 48.73%. This finding aligns with previous studies reporting no significant differences between male and female cattle [23,35], although one study did observe sex-related variation, possibly due to differences in management practices or environmental exposure [25].

Finally, breed had no significant effect on infection rates, with local cattle showing 51.62% prevalence and crossbreds 46.15%. This finding agrees with earlier research [24,34], suggesting similar susceptibility between breeds, although a few studies reported breed-related differences, potentially attributable to genetic factors or breed-specific traits [35,37].

## 5. Limitations

The cross-sectional design and use of a single diagnostic method provide an accurate snapshot of prevalence and associated risk factors within the study population. While longitudinal or multi-method studies may provide additional detail, the current methodology is sufficient to address the study objectives and offers valuable insights for local disease control strategies.

## 6. Conclusion

This study revealed a high overall prevalence of bovine fasciolosis (49.21%) in the Bahir Dar area, confirming its significance as a major parasitic disease affecting cattle in the region. Prevalence varied notably across localities, with Sebatamit exhibiting the highest rate (61.84%) and Latammba the lowest (27.65%). Analysis of risk factors indicated a significant association between body condition and fasciolosis, with poorly conditioned cattle being more susceptible to infection. In contrast, no statistically significant differences were observed based on age, sex, or breed, suggesting that cattle across all demographic groups are at risk. These findings underscore the need for targeted, location-specific control strategies and highlight the importance of improved nutritional and health management practices to reduce the burden of fasciolosis in cattle populations.

## Data sharing statement

Data is provided within the manuscript.

## Ethical Statement

Not applicaple

## Author Contributions

T.M: writing – review & editing, writing – original draft, resources, formal analysis, investigation, conceptualization, methodology, validation. T.S: writing – review & editing, resources, validation, methodology, visualation. A.B.T: writing the original draft, review editing, funding accusation, resources, methodology, conceptualization, formal analysis, supervision and data curation. All authors read and approved the final manuscript.

## Finding

None

### Acknowledgements

The authors extend their sincere gratitude to all the dedicated staff members of the Bahir Dar Regional Veterinary Laboratory.

